# Extracellular Matrix Proteins Changes in the Left Ventricular Unloading and Overloading after Myocardial Infarction

**DOI:** 10.1101/2023.08.28.555160

**Authors:** Spyros A. Mavropoulos, Tomoki Sakata, Renata Mazurek, Anjali J. Ravichandran, Jonas M. Marx, Kiyotake Ishikawa

**Affiliations:** Cardiovascular Research Institute, Icahn School of Medicine at Mount Sinai, New York, NY, USA

**Keywords:** Extracellular matrix, myocardial infarction, left ventricular unloading, proteomics

## Abstract

**Background:** The cardiac extracellular matrix (ECM) is a dynamic scaffold that transmits and responds to forces that act on the heart during the cardiac cycle. While acute left ventricle (LV) unloading by percutaneous LV assist device (pLVAD) is now a therapeutic option, its impact on ECM is unknown. We hypothesize that the composition of LV ECM proteins changes in response to reduced wall stress with mechanical LV unloading.

**Methods:** Ten Yorkshire pigs underwent myocardial infarction, and one week later were subjected to either 2 hours of LV unloading with pLVAD or 2 hours of LV overloading by mechanically-induced acute aortic regurgitation. Non-ischemic remote myocardium was processed for ECM specific proteomics. Volcano plots were constructed to determine differentially expressed proteins between the groups. Gene ontology (GO) analysis was performed to identify associated molecular processes and biological functions. Immunofluorescence microscopy was performed to validate differential protein expression.

**Results:** Of the 986 proteins analyzed, 39 were significantly differentially expressed between overloaded and unloaded myocardium. GO analysis by molecular function revealed that differences tending towards RNA binding and lipid binding proteins. Analysis by biological process showed that cell differentiation, vesicle transport, and programmed cell death were the processes most affected. Staining for three of the identified differentially expressed proteins had results congruent with proteomic analysis.

**Conclusions:** There were significant differences in the protein composition of the cardiac ECM between acutely unloaded and overloaded myocardium, suggesting that it actively responds to altered LV load.

## Introduction

Acute left ventricular (LV) unloading by percutaneous left ventricular assist devices (pLVAD) provides hemodynamic improvement and relieves elevated wall stress by decreasing LV volume and pressure(1,2). It also reduces myocardial oxygen consumption and corrects heart failure-induced derangements in cardiac metabolism(3,4). Benefit of acute LV unloading is now assessed in an ongoing clinical trial investigating whether LV unloading prior to reperfusion can reduce infarct size in acute myocardial infarction patients(5).

The cardiac extracellular matrix (ECM) is a dynamic scaffold that transmits and responds to forces that act on the heart during the cardiac cycle. It is the non-cellular component of cardiac tissue and is composed of glycoproteins, proteoglycans, and glycosaminoglycans that rearrange themselves in response to myocardial injury as part of the remodeling process(6,7). The cardiac ECM modulates a number of processes, including cell signaling, cell adhesion, angiogenesis, and fibrosis, and it has been found that changes in the cardiac ECM can occur as acutely as 12 hours after volume overload(8).

There have been studies that have investigated the intracellular changes that occur during acute LV unloading(9,10), but none so far that examines its effects on the cardiac ECM. Because the cardiac ECM is a dynamic scaffold that both transmits and responds to the forces that act upon the heart during the cardiac cycle, it seems reasonable that the changes in LV pressure and wall stress induced by cardiac unloading would elicit a response from the cardiac ECM. We hypothesize that acute LV unloading with a pLVAD alters the composition of LV ECM proteins in response to reduced wall stress. To better characterize the impact of LV loading condition on the myocardium, unloading group was compared to the overloading group.

## Methods

### Animal Models of Unloading and Overloading

All animal procedures were approved by the Institutional Animal and Use Committee at the Icahn School of Medicine at Mount Sinai, and the care of animals complied with the Guide for the Care and Use of Laboratory Animals (National Institutes of Health publication No. 85-23, revised, 1996). The animals were acclimatized to the facility for at least 72 hours before being enrolled in experiments.

A schematic of the experimental design is shown in Figure 1. Ten Yorkshire pigs (43.6 ± 3.9 kg) were pre-medicated using intramuscular Telazol (tiletamine/zolazepam) (8.0 mg/kg), intubated and ventilated with 100% oxygen. General anesthesia was maintained with intravenous propofol (10mg/kg/h) throughout the procedure. Buprenorphine (0.03 mg/kg) was given prior to the procedure for analgesia. Pigs were subjected to 90-minute balloon occlusion of the middle left anterior descending artery (LAD) and subsequent reperfusion and recovery, as previously described(11). One week after MI induction, the pigs were randomly assigned to either the Unloading group (n=5) or Overloading group (n=5). For the Unloading group, mechanical LV support was initiated using an Impella CP (Abiomed, Danvers, Massachusetts). Fluoroscopy was used to ensure proper positioning of the device and heparin was given intravenously for anticoagulation. Pump support was increased stepwise, and maximal achievable support was maintained in each pig for 2 hours. For the Overloading group, moderate to severe aortic regurgitation was induced by mechanical disruption of the aortic valve using myocardial biopsy forceps. A third group (MI, n=6) of age-matched pigs that only underwent MI induction served as a reference. Animals were euthanized 2 hours after LV unloading or overloading by intravenously injecting the potassium solution (20mEq) and the duplicate 1 cm^3^ samples from the remote non-ischemic myocardium at the pupillary muscle level were collected and immediately frozen for proteomics.

**Figure 1.**
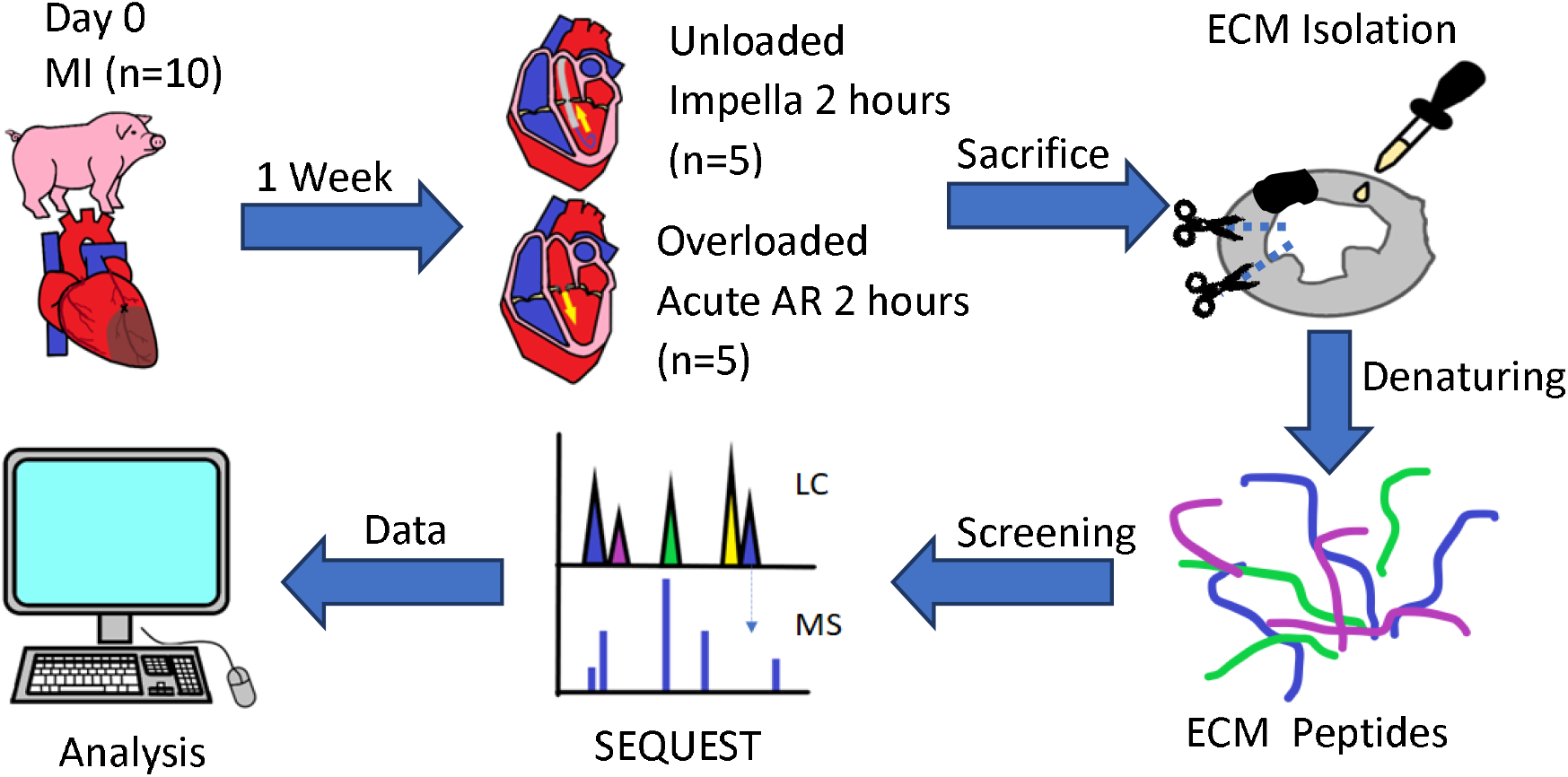
Experimental workflow for the analysis of cardiac ECM proteins in the unloaded and overloaded states.

### ECM Proteomics and Analysis

ECM proteins were isolated and digested from collected myocardial tissue samples as described previously(12). The ECM peptides were then separated by liquid chromatography and identified by mass spectrometry using the SEQUEST library. Volcano plots were constructed by plotting log2(fold-change) vs (-log10(p)) of the resultant data to determine proteins of interest from the data set. Gene ontology (GO) analysis (UniprotKB) was performed to identify associated molecular processes and biological functions of these differentially expressed proteins.

### Immunofluorescence Microscopy

To validate the results of the proteomics analysis, formalin-fixed paraffin embedded tissues sections from remote, non-ischemic myocardium were probed for three of the differentially expressed proteins. The sections were incubated with CAPNS1 (1:500; Biorbyt, Cambridge, United Kingdom), FERMT2 (1:500; Biorbyt, Cambridge, United Kingdom), or APOA4 (1:500; MyBioSource, San Diego, CA) primary antibodies followed by Alexafluor488-conjugated secondary antibody. Nuclei were stained with DAPI. ImageJ (NIH, Bethesda, MD) was used to quantify the intensity of fluorescence for the ECM proteins stained.

### Statistical Analysis

Data are expressed as mean ± standard error of the mean. Groups were compared with one-way ANOVA, with post-hoc Tukey’s multiple comparisons test. All analyses were conducted with Prism version 9.4.1 (GraphPad Software, LLC., San Diego, CA). A p value <0.05 was considered statistically significant.

## Results

In order to increase the sensitivity and screen for as many potential candidates as possible, differential expression was defined as -log10(p) > 1.3 and |log2(fold-change)| > 0.5, which corresponds to a p value < 0.05 and fold change of ± 50%. Using this wide net, of the 986 proteins identified by the SEQUEST library in the cardiac ECM isolates, 39 were identified as significantly differentially expressed between unloading and overloading myocardium (Figure 2).

**Figure 2.**
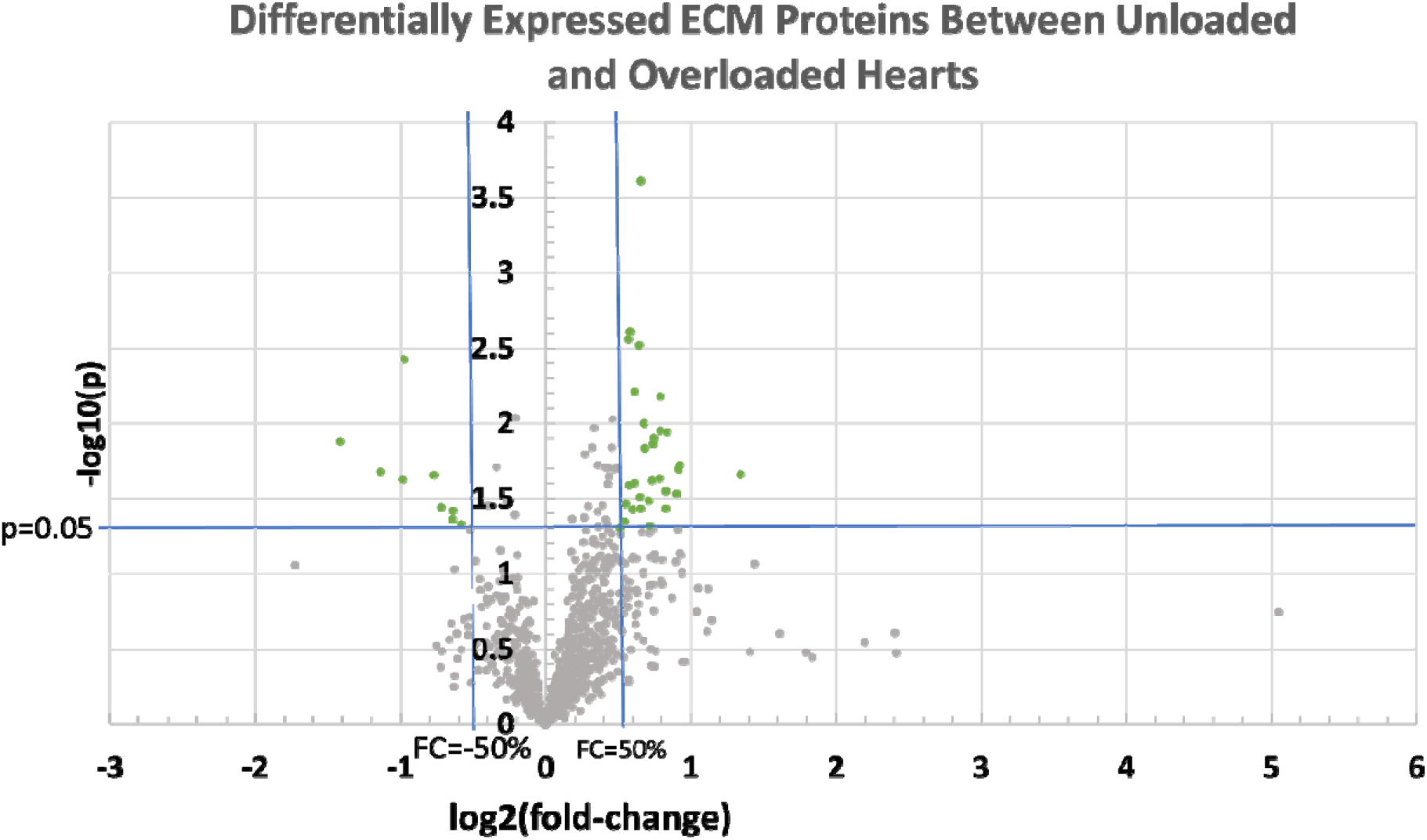
Volcano plot constructed from proteomics data of Unloaded and Overloaded LV ECM isolates. Of the 986 proteins identified by the SEQUEST library in the cardiac ECM isolates, 39 were significantly differentially expressed (-log10(p) > 1.3 and |log2(fold-change)| > 0.5) between loaded and unloaded hearts.

Due to the generally poor annotation for pig genes, GO analysis was performed manually through matching of the human proteins that corresponded to the pig proteins identified by SEQUEST in UniprotKB, and the results are shown in Figure 3. GO analysis of the differentially expressed proteins in the domain of Molecular Function (Figure 3.a) showed that there was favoring of proteins involved in RNA binding (6 proteins) and lipid binding functions (4 proteins). In the domain of Biological Function (Figure 3.b), proteins involved in cell differentiation (7 proteins), programmed cell death (5 proteins), and vesicle transport (5 proteins) had the highest representation.

**Figure 3.**
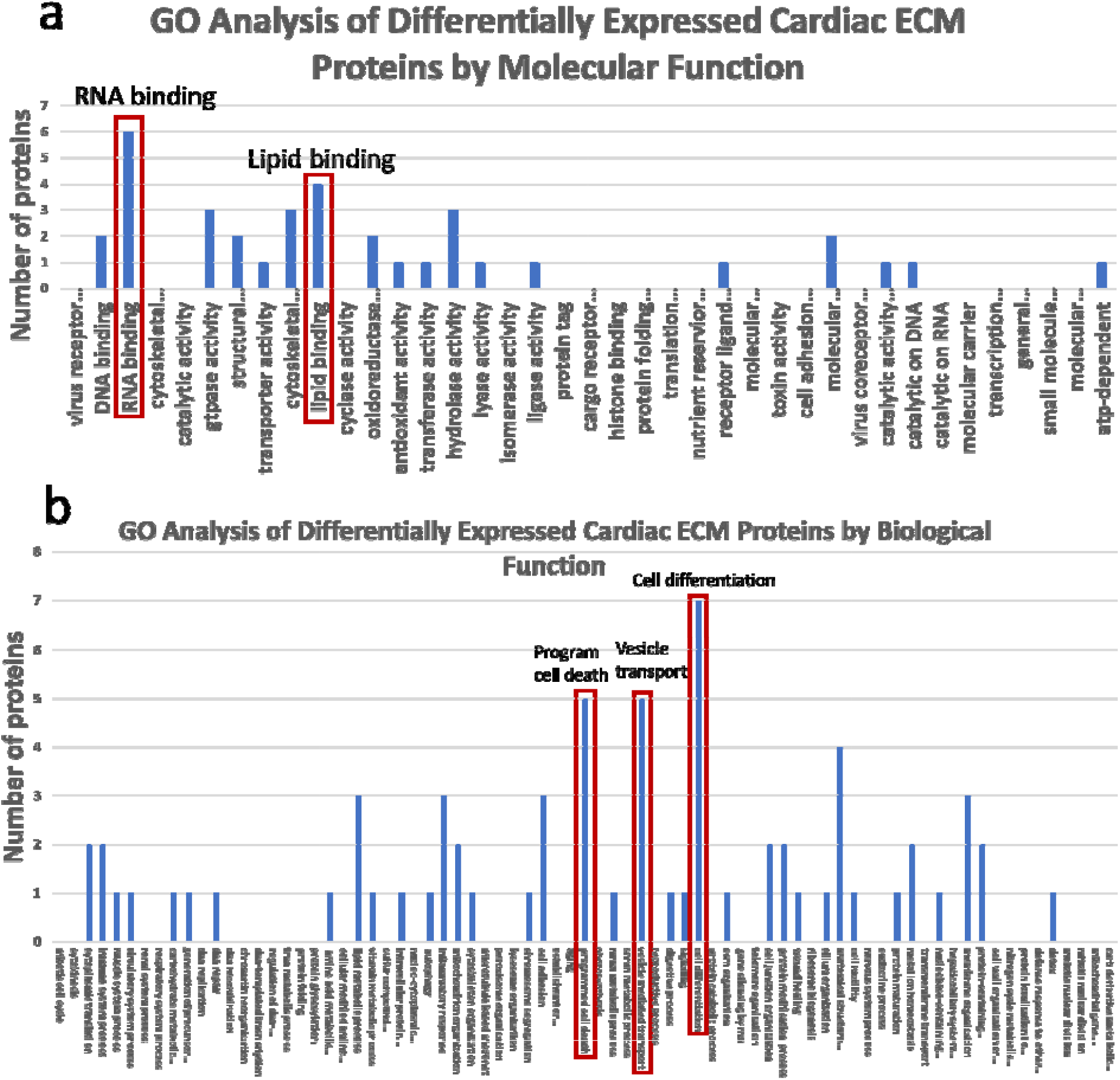
Gene Ontology analysis of the differentially expressed proteins identified. GO analysis of the differentially expressed proteins in the domain of Molecular Function (a) showed that there was favoring of proteins involved in RNA binding and lipid binding functions. In the domain of Biological Function (b), proteins involved in cell differentiation, programmed cell death, and vesicle transport had the highest representation.

The proteins Calpain Small Subunit 1 (CAPNS1), Fermitin Family Homolog 2 (FERMT2), and Apolipoprotein A4 (APOA4) were among the candidates identified as being differentially expressed through proteomic analysis, with significant increases of these proteins in the ECM of unloading hearts when compared to overloading ones. To validate the results of the proteomic analysis, we used immunohistochemistry. Quantitation of the staining intensity exhibited congruent results with those of the proteomics analysis (Figure 4a). For CAPNS1 (Figure 4b), the Unloading group had a mean fluorescence intensity of 1.014 AU compared to 0.1790 AU in the Overloading group (p<0.0001 vs Unloading) and 0.2996 in the MI only group (p<0.001 vs Unloading). For FERMT2 (Figure 4c), mean fluorescence intensity was 0.4329 AU in the Unloading group, 0.01431 AU in the Overloading group (p<0.01 vs Unloaded), and 0.08125 AU in the MI only group (p<0.05 vs Unloaded). For APOA4 (Figure 4d), the Unloading group had a mean fluorescence intensity of 3.327 AU compared to 0.7293 AU in the Overloading group (p<0.05 vs Unloading) and 0.6809 in the MI only group (p<0.05 vs Unloading).

**Figure 4.**
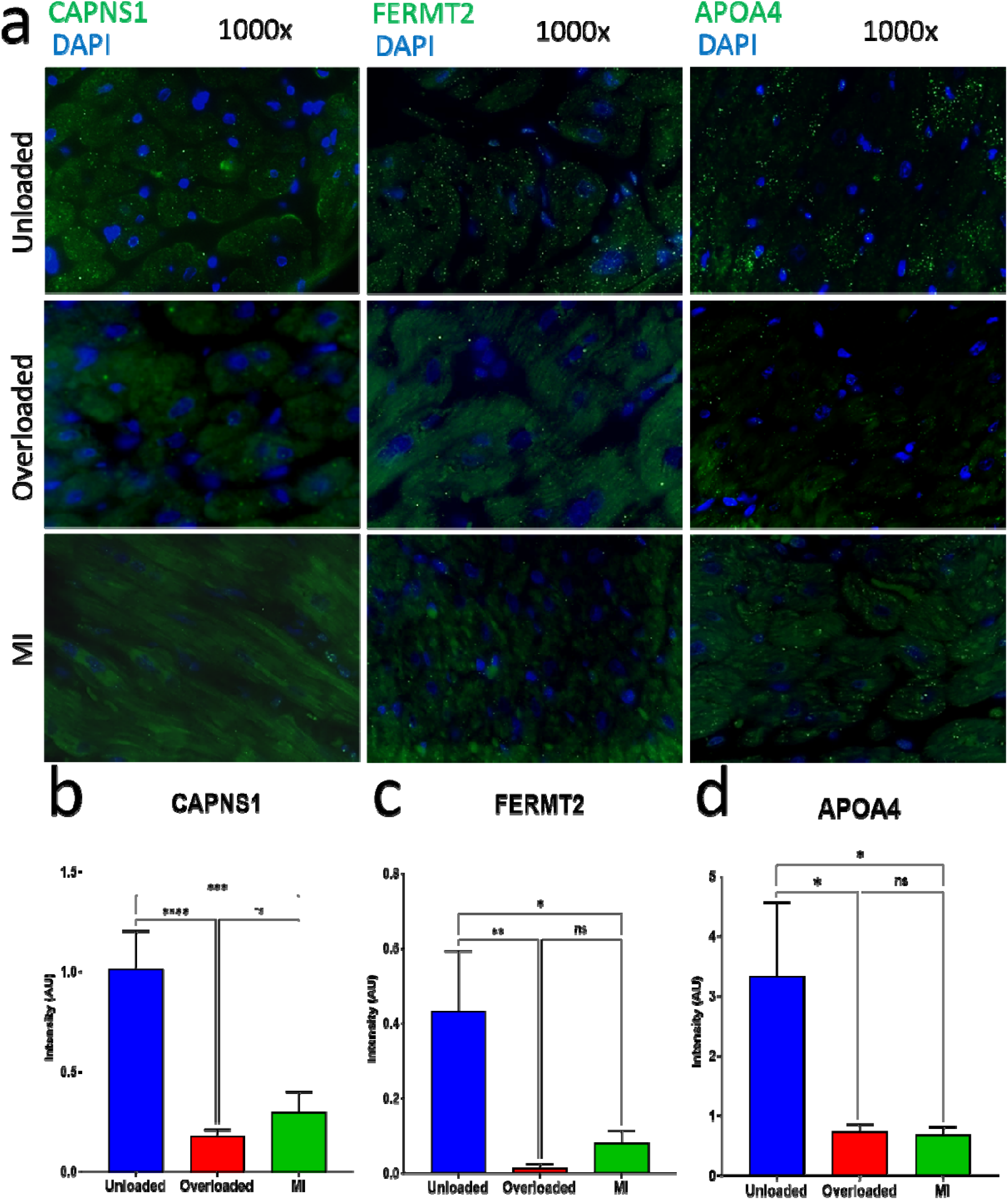
Immunofluorescence microscopy of a few of the differentially expressed proteins identified. CAPNS1, FERMT2, and APOA4 were among the candidates identified as being differentially expressed through proteomic analysis of cardiac ECM. Representative images are shown at 1000x magnification (a). Quantitation of fluorescence intensity of antibodies against CAPNS1 (b), FERMT2 (c), and APOA4 (d) in the LV from an area remote to the infarct area. *p<0.05, **p<0.01, ***p<0.001, ****p<0.0001, ns = not statistically significant.

## Discussion

Since the cardiac ECM both transmits and responds to the forces that act upon the heart, and LV unloading reduces wall stress, it is reasonable to think that the cardiac ECM would respond to the changes in forces that result from LV unloading. Our analysis revealed that there were significant differences in the protein composition of the cardiac ECM between acutely unloaded and overloaded myocardium. It takes approximately 5-10 minutes for transcription induction by a stimulus, and 10-20 minutes for translocation of significant amounts of mRNA to the cytoplasm(13). Due to the acute time period that unloading occurred in this study (2 hours), it is unlikely that these changes were mainly powered by the de novo synthesis of new proteins in response to the changes in LV pressure and wall stress. Rather, it is more likely that either a change in translocation, an increase in trafficking from a remote site, or a change in the rate of degradation of these proteins provided the main contribution to the differential expression of ECM proteins observed between the Unloading and Overloading groups. However, further studies are needed to investigate the mechanism(s) underlying unloading-associated cardiac ECM protein changes.GO analysis revealed that these proteins tended to be involved in RNA binding, lipid binding, cell differentiation, programmed cell death, and vesicle transport. This suggests initiation of changes in cardiac cell phenotype, particularly by those proteins that are involved in RNA binding and those involved in cell differentiation. Whether the changes in the ECM proteome evoked by unloading persist during longer periods of unloading or after the cessation of unloading is still unknown. It is possible that the changes such as cell differentiation induced by pLVAD unloading have benefits that continue long past device removal and should be studied further.

The results of immunofluorescence microscopy of a few of the candidate proteins identified as being differentially expressed through proteomics were in line with what was observed through proteomic analysis. This provides further validation of the quality of the isolation and the results obtained from proteomic analysis of the isolated ECM.

In the future, a study analyzing the potential changes in cardiac ECM due to LV unloading in the setting of heart failure and cardiogenic shock would be important, as they are the population that pLVAD is used as a therapeutic strategy. Such a study would provide further insight into the mechanisms of this therapy for potential benefit other than reduction of oxygen consumption. Investigating whether these ECM changes observed in pig heart tissue can also be seen in human heart tissue would be beneficial. Not only would this further our understanding of the benefits of cardiac unloading, but it would assist in identifying novel potential therapeutic targets in treating heart failure.

### Limitations

This study has several limitations, and the largest one is the unloading/overloading period. A single timepoint of 2 hours was used due to the logistical limits involved in needing to actively anesthetize, sedate, and monitor the pigs for the duration of the unloading period. Therefore, these cardiac ECM proteomics results are merely a snapshot in time. Our data shows that pLVAD unloading can induce changes in the cardiac ECM very acutely, which is certainly important in understanding the impact of unloading on the heart. However, no conclusions can be drawn about the direction or durability of these changes over a longer period of unloading. Patients receiving pLVAD therapy typically have the device for at least a day or longer, so it would be valuable to extend the time frame of the unloading period to 24 hours or longer, to mimic these conditions more accurately.

Another limitation is the lack of inclusion of a group that was unloaded and then recovered. A group where the pLVAD was discontinued and allowed to survive for another 24 hours or longer would be useful in determining whether the cardiac ECM changes observed in this study are enduring and if there are longer-term changes in cell phenotype after acute unloading.

## Conclusions

Acute unloading is capable of evoking rapid changes in the cardiac ECM. There were significant differences in the protein composition of the cardiac ECM between acutely overloaded and unloaded myocardium that suggest initiation of changes in cardiac cell phenotype. It is possible that the changes induced by pLVAD unloading have benefits that continue long past device removal and should be studied further.

